# Reactive and pre-emptive vaccination strategies to control hepatitis E infection in emergency and refugee settings: a modelling study

**DOI:** 10.1101/219154

**Authors:** Ben S Cooper, Lisa J White, Mahveen R Siddiqui

## Abstract

**Background:** Hepatitis E Virus (HEV) is an important cause of morbidity and mortality in emergency and refugee camp settings. Symptomatic infection is associated with case fatality rates of ~20% in pregnant women. However, its epidemiology is poorly understood and the potential impact of immunisation in outbreak settings uncertain. We aimed to estimate key epidemiological parameters for HEV and to evaluate the potential impact of both reactive vaccination (initiated in response to an epidemic) and pre-emptive vaccination.

**Methods:** We analysed data from one of the world’s largest recorded HEV epidemics, which occurred in refugee camps in Uganda (2007-2009), using transmission dynamic models to estimate epidemiological parameters and assess the potential impact of reactive and pre-emptive vaccination strategies.

**Results:** Under baseline assumptions we estimated the basic reproduction number of HEV to range from 3.9 (95% CrI 2.8, 5.4) to 8.9 (5.4, 14.2). Mean latent and infectious periods were estimated to be 34 (28, 39) and 40 (23, 71) days respectively.

Reactive two-dose vaccination of those aged 16-65 years excluding pregnant women (for whom vaccine is not licensed), if initiated after 50 reported cases, led to mean camp-specific reductions in mortality of 10 to 29%. Pre-emptive vaccination with two doses reduced mortality by 35 to 65%. Both strategies were more effective if coverage was extended to groups for whom the vaccine is not currently licensed. For example, two dose pre-emptive vaccination, if extended to include pregnant women, led to mean reductions in mortality of 66 to 82%.

**Conclusions:** HEV has a high transmission potential in refugee camp settings. Substantial reductions in mortality through vaccination are expected, even if used reactively. There is potential for greater impact if vaccine safety and effectiveness can be established in pregnant women.

**Funding:** Wellcome Trust (106491/Z/14/Z and 089275/Z/09/Z). BC: MRC/DfID (MR/K006924/1).

## Introduction

Communicable diseases are responsible for large excess mortality and morbidity in complex emergencies.^1^ Epidemic Hepatitis E is a particular concern due to high mortality in pregnant women and lack of interventions of proven effectiveness in emergency settings. ^2^ Hepatitis E is caused by a single-stranded RNA virus from the Hepeviridae family. There is one serotype but four genotypes.^3^ Hepatitis E virus (HEV) is enterically-transmitted and has caused large outbreaks in many regions including China, the Indian subcontinent, central Asia and East Africa;^4^ globally, it is a leading cause of acute viral hepatitis and has been estimated to account for ~10,000 annual pregnancy-related deaths in southern Asia alone.^5^ Genotypes 1 and 2 dominate in Africa and most of Asia and are found only in humans; they have been estimated to cause over 20 million infections annually in these regions.^6^ Recently, large outbreaks associated with high mortality have occurred in internally-displaced persons (IDP) camps in Sudan, Uganda and South Sudan.^7–10^.

The risk of symptomatic illness given HEV infection is thought to increase with age and has been estimated to be about 20% in adults infected with genotypes 1 or 2; ^6^ the probability of death given symptomatic infection has been estimated to be approximately 2% for non-pregnant cases and 20% in pregnant women.^6,11^.

With over 60 million people living as refugees or otherwise forcibly displaced worldwide,^12^ the majority in areas vulnerable to HEV infection, understanding HEV epidemiology and the potential for its control is of considerable importance.

## Course of hepatitis E infection

Knowledge of the course of HEV infection is based mainly on two volunteer studies ^13,14^ and four patient studies.^15–17^ The two single volunteer studies (with known dates of exposure) gave incubation periods of 38 and 39 days respectively.^13,14^ Clinical illness typically lasts 1-4 weeks.^17^ Viral shedding in the stool begins about 2 weeks after infection and continues for up to four to five weeks.^18-20^ The start of the infectious period appears to approximately coincide with the onset of the prodromal phase of disease.^14^

## Vaccine

There is a recombinant vaccine, HEV 239, which is approved for use in China in those aged 16–65 years who are not pregnant.^21^ It is produced with a genotype 1 isolate and efficacy against both genotypes 1 and 4 has been established in non-human primates. The vaccine has been demonstrated to have >90% efficacy based on a clinical trial involving 109,959 people at risk of HEV infection in an endemic setting, primarily with genotype 4.^21^ Preliminary observations suggest the vaccine is also safe and effective in pregnant women.^22^ Clinical trial data are lacking for those aged <16 or >65 years, in areas where genotypes 1 and 2 dominate, and in outbreak settings.

We aimed to quantify key epidemiological parameters for HEV in refugee camp settings and evaluate the potential benefits of vaccination. We consider both pre-emptive (prior to HEV cases occurring) and reactive vaccination (once HEV outbreaks are already underway), and evaluate the potential impact of selecting different target groups to receive the vaccine.

To do this we fitted dynamic transmission models to data from three large HEV epidemics in refugee camps. We used a Bayesian framework to combine data from previous studies with observed epidemic data to obtain an improved understanding of the natural history of HEV infection, quantify the transmission potential, and evaluate the potential for vaccination to reduce the number of clinical cases and associated mortality.

## Methods

### Data

Data came from three epidemics in 2007-2009 from internally displaced persons (IDP) camps in the district of Kitgum, Uganda: Agoro, Madi Opei, and Paloga (estimated populations 16,689, 10,442, and 10,555 respectively). Jaundice cases were recorded in facility-based passive surveillance systems. Evidence from serology and reverse transcription–PCR confirmed HEV to be the outbreak cause; other causes of viral hepatitis were rare.^10^.

### Transmission model

We fit a series of deterministic transmission models to the data, assuming latent and infectious periods were common to all camps but allowing transmissibility to vary by camp. In our baseline model individuals were assumed to be in one of four possible states: susceptible to infection (S); latently-infected but not yet infectious (E); infectious (I); and recovered and immune (R). The rate at which susceptibles became infected was assumed to scale linearly with the number currently infectious. Information from previous studies was used to construct moderately informative prior distributions (*priors*) for natural history parameters. These priors represent knowledge about disease progression parameters before fitting the model to the outbreak data. When combined with analysis of the data they give rise to posterior distributions, which represent what we know about the parameters after analysing the new data.

A human challenge study had previously reported that viable virus could be found in the faeces four days before onset of the icteric phase of disease.^14^ We therefore assumed that patients become infectious one week before the icteric phase of disease. A prior for the mean infectious period was derived from a study of faecal shedding in 11 patients with sporadic acute HEV infection acquired in Bangladesh, Vietnam, Nepal, and Japan (see Supplementary material Table S1);^20^ this study found that viable HEV could be recovered from faecal samples up until 2-5 weeks after hepatitis onset. The prior distribution for the latent period was based on a single observation from the same study where there was a delay of 34 days between inoculation and viable HEV in faeces. Faecal shedding of HEV in asymptomatically infected people is known to occur;^23^ we assumed no difference in faecal viral shedding between symptomatic and asymptomatic individuals. We used seroprevalence data to derive an informative prior for the proportion of infections that are reported (see Supplementary material, Table S1). We assumed no immunity to HEV in the refugee camp populations prior to first reported case, consistent with the absence of reports of previous HEV epidemics in Kitgum district and serological data.^10^ We performed multiple sensitivity analyses, considering models with: i) different assumptions about the distribution of the latent and infectious periods; and ii) a camp-specific transmission rate affected by a water and sanitation intervention. All the above models assumed nothing about the relative importance of different modes of transmission and are consistent with both transmission mediated by a locally contaminated environment (though without long-term virus persistence in this environment) and direct person-to-person spread. We also compared our findings with those from a model explicitly accounting for a second mode of transmission, an unobserved environmental reservoir of HEV where virus may persist for longer time periods. We performed an extensive sensitivity analysis by running this model under 25 different prior assumptions representing combinations of five different assumptions about the relative importance of this environmental reservoir in the early stages of an epidemic and five different assumptions about persistence of viable virus in the environment.

### Intervention analysis

To evaluate the potential impact of vaccination we used estimates of vaccine effectiveness after 2 and 3 doses derived from data in Zhu et al ^21^ (see Supplementary material) and assumed 90% coverage for the first two doses in target groups. The intervals between the first and second and the second and third doses were one and five months respectively. There was no evidence of any effect of a single dose of vaccine in the clinical trial, so this was excluded from the analysis. In the absence of evidence to the contrary, we assumed vaccine effectiveness did not vary by age.

Case fatality ratios amongst those pregnant and those not pregnant were derived from the meta-analysis of Rein et al.^6^ We assumed a threshold of 50 or 100 reported cases as the starting point for reactive vaccination. Other assumptions are listed in Table 1.

In all simulations we accounted for uncertainty in model parameters by drawing these from the posterior distributions obtained by fitting the model to data from the three camps.

### Inference and simulation details

The posterior distributions for model parameters were obtained by fitting the models to data using a Markov chain Monte Carlo approach assuming that observed weekly cases followed a negative binomial distribution with a mean given by the deterministic model. Analysis was performed in R.^24^ Further technical details can be found in the Supplementary material.

## Results

The baseline model gave good fits to the three epidemic curves, with the observed number of weekly cases usually within the 95% prediction intervals (Figure 1).

**Figure 1.**
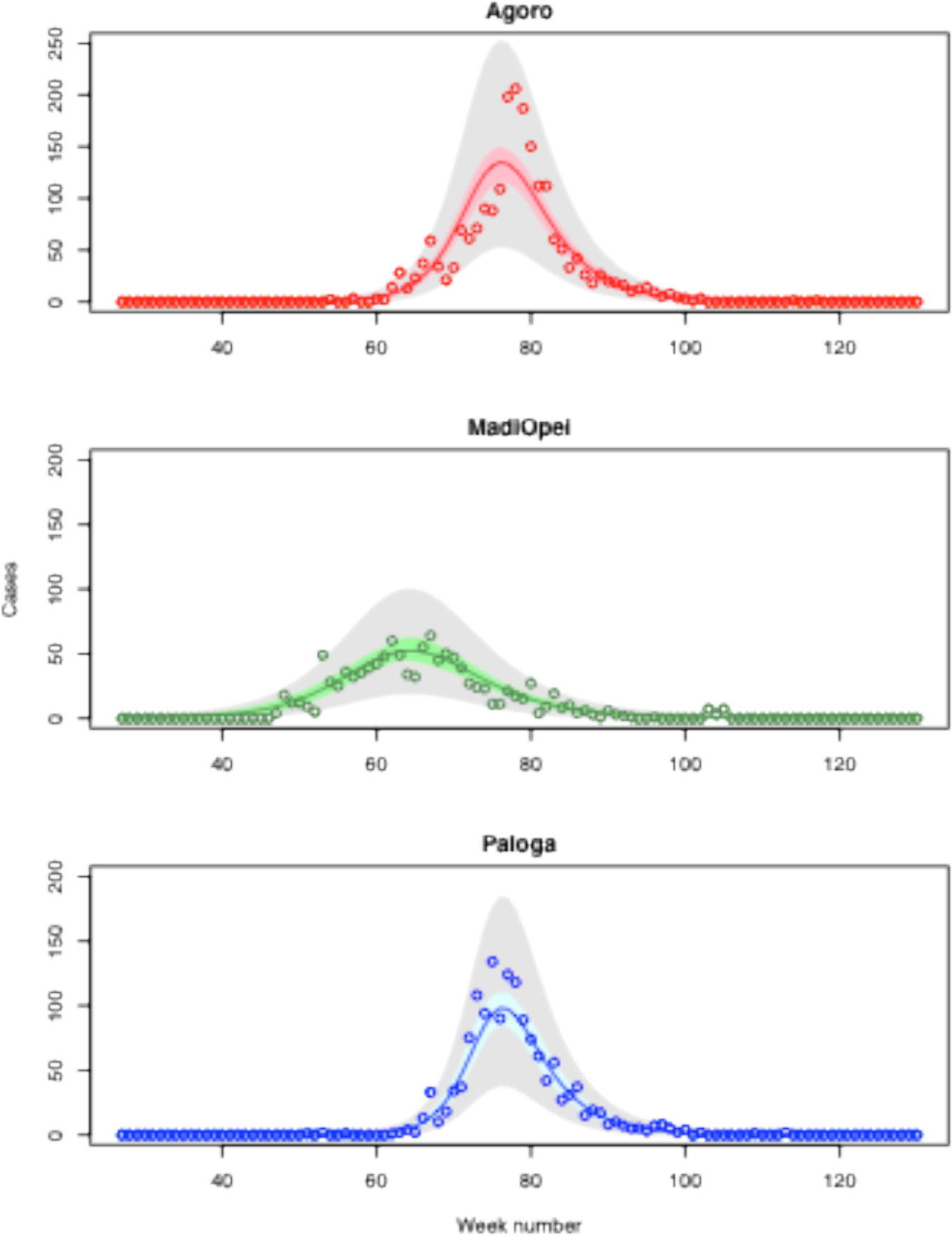
Observed and predicted hepatitis E cases. Observed weekly cases (circles) and expected weekly cases at the three sites based on the baseline transmission model with moderately informative priors (solid line) and 95% Credible Intervals for the mean (shaded coloured region) and for the predicted number of cases (shaded grey region). Week number 1 corresponds to the first week of 2007.

Under the baseline model the mean latent period was estimated to be 34 days (95% Credible Interval (CrI): [29, 39]) (Figure 2 and Supplementary material Table S3). Similar estimates were obtained in sensitivity analyses, though a shorter mean latent period (19 days [10, 34]) was inferred if alternative distributional assumptions were made about the latent period (Model 3, Supplementary material Table S3). The mean infectious period was estimated to be 36 days, 95% CrI (21, 64), in the baseline model, though this reduced to 27 days (21, 37) under different distributional assumptions (Model 4, Supplementary material Table S3). The estimated proportion of infections reported was 12.5% (11.4%, 13.6%) in the baseline model and similar in all sensitivity analyses. Under baseline assumptions the basic reproduction numbers were estimated to be similar in two of the three camps (Agoro 6.5 (4.5, 9.9); Paloga 8.5 (5.3, 11.4)), but smaller in Madi Opei (3.7 (2.8, 5.1)). Central estimates for these reproduction numbers were similar in most sensitivity analyses (Supplementary material Table S3), though higher in models that explicitly accounted for two modes of transmission (Supplementary material Table S4).

**Figure 2.**
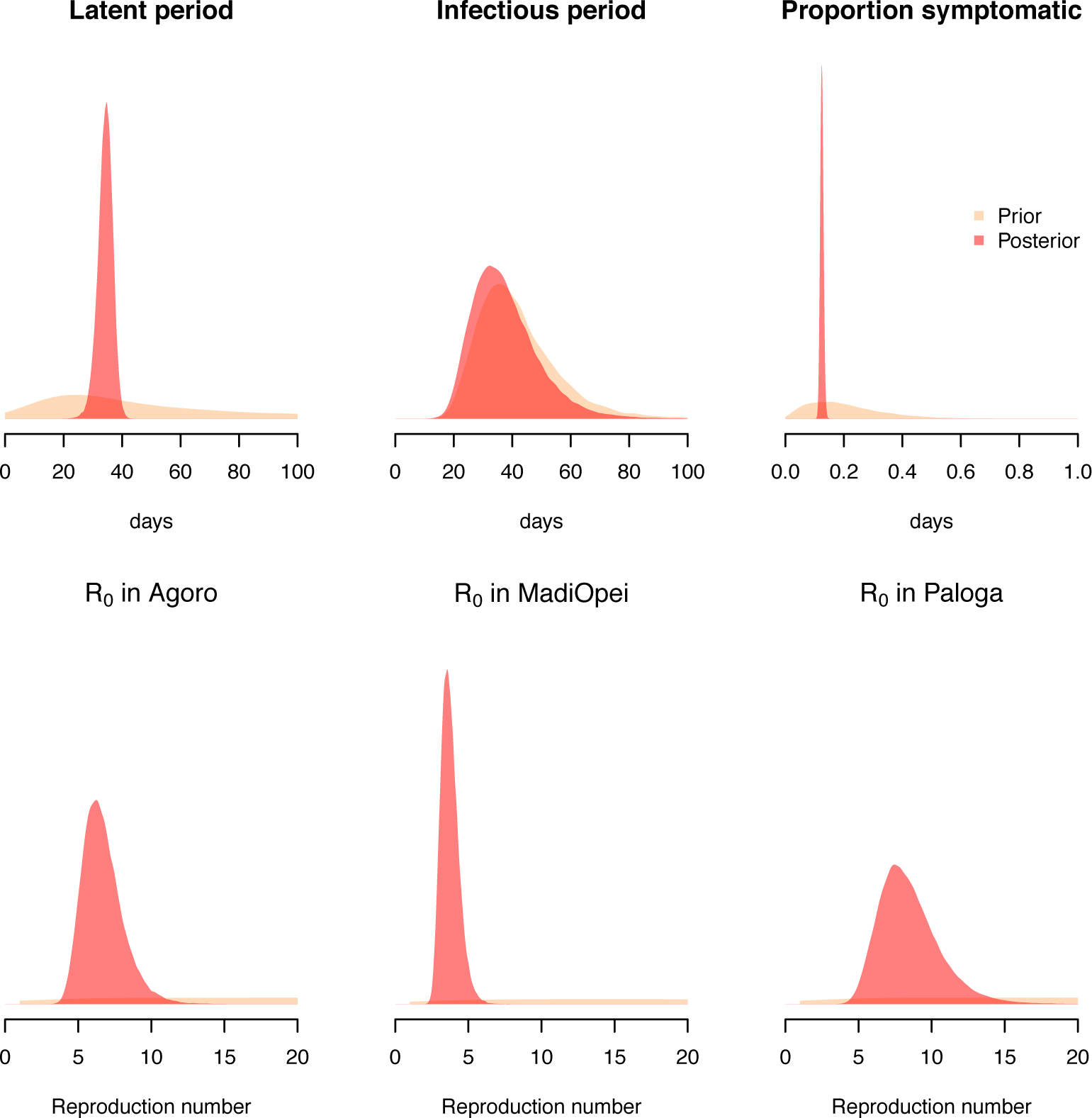
Prior and posterior distributions for key epidemiological parameters. Estimates are derived using the baseline SEIR model with moderately informative priors: the mean latent period, the mean infectious period and the proportion symptomatic (top row) are assumed to share the same distributions at the three camps. Posterior distributions of the basic reproduction number (R_0_) are allowed to vary by camp (bottom row).

Analysis of the data using models explicitly accounting for the water and sanitation intervention (models 2a and 2b, Supplementary material Table S3) did not provide evidence that these interventions were effective in reducing transmission. However, results were consistent with both substantial positive and negative effects, indicating that the data contained little information about the effects of the water and sanitation responses on transmission. This reflects the fact that in all three camps the interventions were not fully in place until the HEV epidemics were almost over (Supplementary material Figure S1). Fitting the models accounting for a second mode of transmission corresponding to an unobserved environmental reservoir showed that the data were only consistent with the majority of transmission occurring via this route if mean persistence of viable virus in this environmental reservoir was two weeks or more (Supplementary material Figure S4).

Considering the potential effects of different vaccine usage scenarios, under the baseline model, reactive vaccination (assuming two doses after the first 100 or 50 cases) was capable of producing important though relatively modest reductions in cases and deaths (Figure 3). These reductions were sensitive to the threshold number of cases before reactive vaccination was initiated: when the threshold was 100, reductions in mortality compared to no vaccination scenarios were unlikely to exceed 25% even assuming the vaccine was given without age or pregnancy restrictions; for a threshold of 50, mortality reductions of about 40% were plausible. In contrast, with no age-restriction on vaccine recipients and pre-emptive vaccination, reductions in mortality of 100% were possible indicating that the herd immunity threshold had been reached. This was true whether two or three doses were administered, though in the former case uncertainty was far larger reflecting the lower precision in the estimated vaccine effectiveness of two doses.

**Figure 3.**
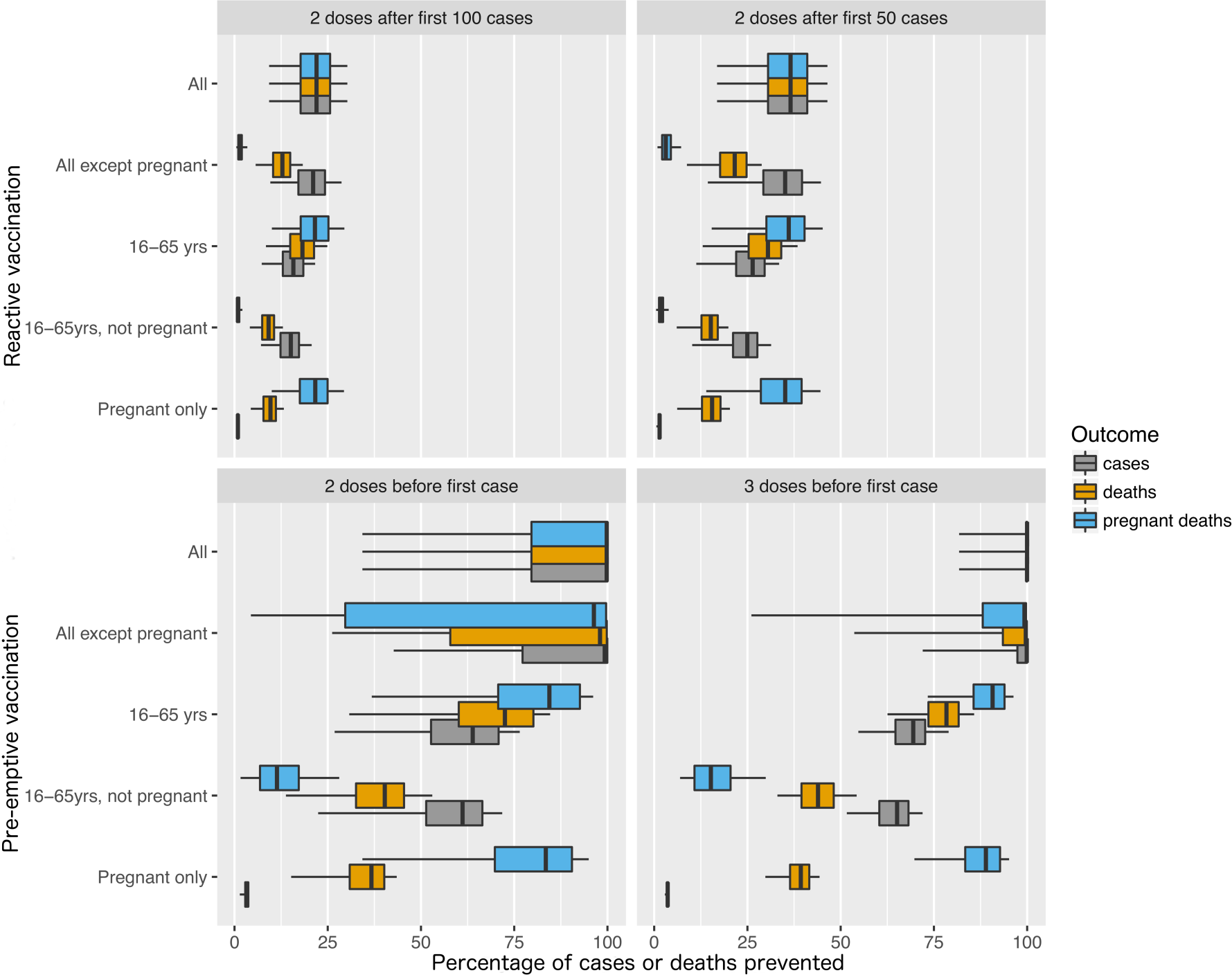
Percentage of hepatitis E cases, deaths and deaths of pregnant women prevented by different vaccination strategies. Estimates are derived using posterior distributions of model parameters obtained by fitting the baseline model with moderately informative priors to data from Agoro. Boxplots show the median (central bar), interquartile range (extent of coloured box) and 5th and 95th percentiles (whiskers).

The impact of excluding pregnant women from the population to vaccinate varied with scenario. In cases where vaccination had a high chance of achieving herd immunity (i.e. when vaccine was used pre-emptively without age restriction) excluding pregnant women had only a moderate negative impact on reductions in mortality. In contrast, when herd immunity was less likely to be obtained through vaccination (i.e. when the vaccine was used reactively or with age restrictions) excluding pregnant women led to substantially smaller reductions in total mortality (and much smaller reductions in mortality in pregnant women).

Comparing vaccination policies with and without age restrictions, restricting receipt of the vaccine to those between the ages of 16 and 65 years had a small impact on reductions in mortality in the reactive vaccination strategy, but a far larger negative impact on the pre-emptive vaccination strategy, a consequence of reducing the chance of achieving herd immunity in these latter strategies.

These broad conclusions about the impact of different vaccination strategies were robust to precise details of model specification. In particular, under all 25 scenarios in the model with two modes of transmission, similar patterns were seen (Supplementary material Figure S5), though the smallest reductions in mortality were seen when priors expressed the belief that most transmission was via this environmental route and viral persistence in this environment was long.

## Discussion

Our results indicate that in refugee camp settings HEV can be highly transmissible. In one camp (Paloga) the estimated mean number of secondary cases per primary case at the start of the epidemic (the basic reproduction number, *R_0_*,) exceeded 6.5 under all model assumptions. This number has important implications for vaccination policies. To achieve herd immunity requires successful immunization of a percentage of individuals given by 100-100/ *R_0_*.^25^ Thus, even taking the optimistic *R*_0_ value of 6.5, we would require 85% of the population to be effectively immunized to achieve herd immunity. Herd immunity is desirable as it means a major epidemic will not be possible even though many in the population remain susceptible to infection. It is of particular relevance here because HEV-infected pregnant women face a greatly increased risk of mortality but the safety and efficacy of the vaccine in pregnant women remains to be established.

Even when restricting vaccine use to non-pregnant 16-65 year olds (for which the vaccine is licenced in China), the benefits could be substantial, with reductions in mortality of over 40% likely under baseline assumptions given pre-emptive vaccination with three doses. A robust finding was that pre-emptive vaccination was much more effective at preventing HEV cases and deaths than reactive vaccination.

Our work sheds light on other important aspects of HEV epidemiology: we found consistent evidence that the mean latent period is between about 20 and 40 days and that a little over 10% of individuals infected with HEV are identified as cases. The data were less informative about the mean infectious period, though were consistent with the range 20-70 days suggested by previous data.

Other aspects of the epidemiology are less clear. Transmission of HEV is widely thought to occur predominantly via the faecal-oral route, usually through contact with contaminated water; it might therefore be assumed that household transmission is rare. However, a case-control study of 112 symptomatic cases and 145 controls in Paloga found only two behavioural risks associated with symptomatic HEV infection (with adjusted odds ratios of 3 and 2 respectively): use of wide-mouthed water storage vessels and communal hand washing.^26^ Based on these findings, and the presence of HEV RNA in hand-rinse samples, the authors concluded that water storage practices could have played an important role and that transmission was likely to have included household-level spread. Our findings add further weight to these observations. They show that the epidemic curves in all three camps can be reproduced without positing the existence of a saturating environmental reservoir (and models with such a reservoir did not improve fits to data). We found no evidence that the water and sanitation interventions reduced transmission, though these interventions were introduced late and the data provide little evidence for or against their effectiveness. Previous modelling has shown that such interventions have the potential to be highly effective if assumed to be capable of interrupting the dominant mode of transmission.^27^

Given the uncertainties about transmission routes and fundamental methodological challenges in making inferences about an unobserved environmental reservoir,^28^ we performed extensive sensitivity analyses. While this did not make it possible to quantify the relative importance of different transmission routes it did shed light on the circumstances where such a reservoir would be consistent with the observed epidemic data. A key finding is that if the mean persistence of viable virus in the environment was seven days or fewer, the saturating environmental reservoir was estimated to play only a minor role in the epidemic. If mean persistence was two weeks or more, however, the data were compatible with such an environmental reservoir representing the dominant mode of transmission.

Strengths of our work include the use of high quality epidemic data, extensive sensitivity analysis, and an analysis that allows us to incorporate information from previous HEV investigations. This work also has important limitations. First, estimates of HEV vaccine effectiveness from a trial in China may not generalise to a typical emergency setting or to age/ethnic groups not included in the trial. Second, our analysis neglects age, spatial and household structuring, behavioural, biological and temporal heterogeneities that might affect HEV transmission. These may all play important roles in HEV epidemiology in emergency settings but we lacked data of sufficient resolution to meaningfully incorporate them. Delineating such factors is an important area for future work. We assumed no prior immunity though lacked pre-epidemic sera to enable us to assess this assumption. If some people were immune prior to the epidemic, we are likely to have underestimated the basic reproduction numbers. Finally, lacking any data on the efficacy of a single vaccine dose, we assumed it conferred no protective effect. This is a conservative assumption and may mean that we have underestimated the potential vaccine benefits.

In conclusion, this work has shown a high transmission potential for HEV in refugee camp settings, shed light on important natural history parameters, and has shown that mass vaccination campaigns in such high risk populations have the potential to lead to substantial reductions in mortality.

In 2015 the Strategic Advisory Group of Experts on immunization could not recommend the routine use of the vaccine for population sub-groups including children aged less than 16 years and pregnant women but emphasized that the use of the vaccine during outbreaks of hepatitis E should be considered.^29^ Our findings show that such an intervention could have a major impact, particularly if pre-emptive vaccination is used and vaccination can be safely extended to high risk groups excluded from vaccine trials. In particular, these results underline the need to prioritise evaluations of the vaccine in pregnant women.^30^

## Acknowledgements

Geoff N. Mercer (deceased 12th April, 2014).

## Author contributions

BSC developed and implemented the models, analysed the data, performed the literature search and produced the figures. MRS came up with the research questions, curated the data, analysed the data and searched the literature. BSC, MRS and LJW contributed to the design of the study, development of models, interpretation of results and writing the manuscript.

## Ethics committee approval

Ethics committee approval was not required for this study.

